# Sexually Dimorphic Transdifferentiation of Endothelial Cells in Atherosclerosis

**DOI:** 10.64898/2025.12.01.691732

**Authors:** Cheng Luo, Songmao Wang, Yuyan Lyu, Qing Rex Lyu

## Abstract

**Objective:** To elucidate the sex-specific trajectories of endothelial cell (EC) transdifferentiation and their contribution to distinct atherosclerotic plaque phenotypes in males and females.

**Approach and Results:** We utilized *Cdh5-CreER^T^*^2^*/Apoe^−/−^/Rosa26-mTmG* lineage-tracing mice to perform single-nucleus RNA sequencing (snRNA-seq) of aortic tissues from male, female, and ovariectomized female mice following 12 weeks of high-fat diet. We mapped high-resolution EC fates, identifying distinct transdifferentiation routes into smooth muscle cell (SMC)-like, adipocyte-like, macrophage-like, and fibroblast-like cells. Females exhibited higher overall EC plasticity with a preference for SMC-like and metabolically active adipocyte-like trajectories, contributing to plaque stability. Conversely, male ECs favored inflammatory macrophage-like and stress-responsive fates. Trajectory analysis revealed a novel direct EC-to-adipocyte-like transition that bypasses the canonical endothelial-to-mesenchymal transition (EndoMT). Gene regulatory network analysis, validated by in vitro knockdown assays, identified KLF2 as a homeostatic brake on transdifferentiation, while CREB5 and ESRRG emerged as critical molecular switches governing the bifurcation between adipogenic and myogenic fates. Ovariectomy in females shifted the transcriptomic landscape toward a male-like, lipid-anabolic phenotype.

**Conclusions:** EC plasticity is a sex-dimorphic process driving differential plaque composition. We identify a novel, direct EC-to-adipocyte trajectory and pinpoint KLF2 and CREB5 as key regulators, offering new mechanistic insights into why premenopausal women develop more stable atherosclerotic lesions than men.

## Introduction

Atherosclerosis is the primary underlying cause of cardiovascular diseases (CVDs), leading to myocardial infarction (MI) and ischemic stroke ^1,2^. The pathogenesis of atherosclerosis is multifactorial, driven by elevated low-density lipoprotein cholesterol (LDL-C), triglyceride-rich lipoproteins, disturbed hemodynamics, and endothelial barrier dysfunction ^3^. Although lipid-lowering therapies are widely standardized, the global burden of atherosclerosis and its complications continues to rise ^4,5^, highlighting persistent gaps between current mechanistic knowledge and therapeutic efficacy. Therefore, elucidating the cellular and molecular drivers governing the initiation and progression of atherosclerosis remains an urgent priority.

Epidemiological and clinical data have long established prominent sex differences in atherosclerosis ^6–8^. Premenopausal women benefit from the protective effects of endogenous estrogens, typically presenting with lower plaque burden, smaller lipid cores, reduced inflammation, and thicker fibrous caps, features consistent with stable lesions ^9^. In contrast, men typically exhibit higher atherosclerotic incidence and greater plaque burden, characterized by extensive inflammatory infiltration, large lipid rich necrotic cores, and thin fibrous caps, which collectively increase plaque vulnerability and cardiovascular risk ^10^.

This hormonal protection diminishes post-menopause, as declining estrogen levels correlate with a marked increase in cardiovascular risk and accelerated disease progression ^10^. Despite these well documented clinical disparities, mechanistic insights into how sex shapes cellular fate during atherogenesis remain limited, hindering the development of sex specific therapeutic strategies.

Endothelial cells (ECs) form a monolayer lining the luminal surface of the vasculature, serving as a dynamic interface between circulating blood and the underlying smooth muscle cells (SMCs) ^11,12^. Disruption of this barrier allows immune cell infiltration into the subintimal space, triggering inflammatory signaling and driving SMCs to dedifferentiate from a contractile to a synthetic phenotype, characterized by increased proliferation, migration, and cytokine production ^13^. ECs also express mechanosensory molecules that respond to changes in blood flow; sensing of disturbed or oscillatory flow promotes endothelial dysfunction and the development of vascular disease ^14,15^. Furthermore, ECs release vasoactive mediators that regulate vascular tone ^16^, and undergo endothelial-to-mesenchymal transition (EndoMT) under both physiological and pathological conditions, contributing to neointimal expansion, extracellular matrix remodeling, and plaque instability ^17–19^. Collectively, these processes position EC dysfunction as a central contributor to atherogenesis ^20,21^. However, whether and how EC plasticity and transdifferentiation trajectories diverge between sexes to drive distinct plaque phenotypes remains poorly understood.

Lineage tracing serves as a powerful tool to define cell fate during dynamic physiological or pathological processes in vivo ^22^. By crossing tissue specific recombinase mouse lines with reporter strains, the progeny of specific cell types can be permanently labeled and tracked^23^. While EndoMT is a well characterized EC fate in cardiac development and fibrosis ^24^, the full spectrum of EC plasticity in atherosclerosis, particularly regarding sex-specific differentiation routes—has not been comprehensively mapped ^11,24^. In this study, we employed *Cdh5-CreER^T^*^2^*/Apoe^−/−^/Rosa26-mTmG* mice to perform single-cell lineage tracing during atherosclerotic plaque formation in both sexes. These analyses provide a high-resolution view of endothelial dynamics, revealing distinct, sex dependent transdifferentiation trajectories that may explain the differential vascular phenotypes observed in males and females.

## Materials and Methods

### Mouse Line and Atherosclerosis Model Induction

All animal protocols and procedures were approved by the Animal Research Committee of Yishang Biotech Co., Ltd. (Shanghai, China) (IACUC-2024-Mi-006) and conducted in accordance with National Institutes of Health Guidelines on the Care and Use of Laboratory Animals (NIH Publication, 8th Edition, 2011). *Cdh5-CreER^T^*^2^*/Apoe^−/−^/Rosa26-mTmG* mice were generated by breeding *Cdh5-CreER^T^*^2^ (#C001330, Cyagen, China), *Apoe^−/−^* (#C001067, Cyagen, China), and *Rosa26-mTmG* (#C001192, Cyagen, China) mice. In total, 20 male and 25 female *Cdh5-CreER^T^*^2^*/Apoe^−/−^/Rosa26-mTmG* mice were used, of which 5 females underwent ovariectomy. To induce Cre activity, 75 mg/kg/d of tamoxifen (#HY-13757A, MedChemExpress, USA) was injected intraperitoneally for five consecutive days. Mice were fed a high-fat diet (#D12109, ResearchDiets, USA) from 8 weeks of age for a continuous 12 weeks to induce atherosclerotic plaque formation. The successful model establishment was determined by *en face* oil red O staining of the aorta from a random mouse.

### Immunofluorescence Staining

Paraffin-embedded sections (3-5 μm thick) were deparaffinized, rehydrated, and subjected to antigen retrieval. Then, sections were incubated with primary antibodies against CD31-488 (1:200, #CL488-65058, Proteintech, USA) and COL8A1 (1:200, #17251-1-AP, Proteintech, USA), FMO2 (1:200, #67019-1-lg, Proteintech, USA), Ki67(1:100, #AB2008, Beyotime, China), Ly6A (1:200, #14-5981-82, Invitrogen, USA), Myh6 (1:00, #AF7530, Beyotime, China), SMMHC (1:200, #21404-1-AP, Proteintech, USA), and SOX17 (1:200, #NBP2-24568, Novus, USA), overnight at 4°C in the dark. After PBS washing for three times, Alexa Fluor 594-conjugated secondary antibodies (1:200, Invitrogen, USA) were applied for 1 h at 37°C in the dark. Sections were mounted with ProLong Gold anti-fade reagent with 4′,6-diamidino-2-phenylindol (DAPI) (#P36931, Invitrogen, USA) for confocal microscopy (Olympus, Japan).

### Cell Culture and siRNA Transfection

The immortalized human aortic endothelial cell line, TeloHAEC, was purchased from American Type Culture Collection (#CRL-4052, ATCC, USA). Cells were grown in Vascular Cell Basal Medium (#PCS-100-030, ATCC, USA) supplemented with Vascular Endothelial Cell Growth Kit-VEGF (#PCS-100-041, ATCC, USA), and cultured in a 5% CO_2_ humidified incubator at 37°C.

Small interfering RNAs (siRNAs) targeting *KLF2* and *CREB5* were synthesized by SYNBIO TECHNOLOGIES (Suzhou, China). The siRNA sequences for *KLF2* are: siKLF2-#1, CGGCACCGACGACGACCUCAA; siKLF2-#2, GGUAUUUAUUGGACCCAGA; siKLF2-#3, CGCCUUUCGGUGGCCCUGGUU. The siRNA sequences for *CREB5* are: siCREB5-#1, ACAUGCAGCUUCAGAAUGA; siCREB5-#2, ACAUGCAGCUUCAGAAUGA; siCREB5-#3, GAAUCACAAGGAUAUCUAA. All siRNAs were transfected using Lipofectamine™ RNAiMAX (#13778150, Invitrogen, USA) according to the manufacturer’s instructions. Briefly, siRNA and RNAiMAX were each diluted in Opti-MEM™ (#31985070, Thermo Fisher Scientific, USA), mixed gently, incubated at room temperature for 15 min, and then added to cells at a final siRNA concentration of 50 nM. The culture medium was replaced with fresh growth medium after 24 hrs.

### RNA Purification and RNA Sequencing

Total RNA was extracted using TRIzol reagent (#15596026, Thermo Fisher Scientific, USA) following the manufacturer’s instructions. RNA sequencing was performed by Shanghai OEBiotech Co., Ltd. (Shanghai, China). Paired-end sequencing was conducted on an Illumina NovaSeq™ 6000 platform (Illumina, USA). Differentially expressed genes (DEGs) were identified using the *DESeq2* package with default parameters. Genes with an adjusted *p*-value < 0.05 and |log₂(Fold Change)| ≥ 1 were considered significantly differentially expressed and subsequently used for Gene Ontology (GO) and Kyoto Encyclopedia of Genes and Genomes (KEGG) enrichment analyses.

### Nucleus Isolation of the Aortic Tissue

The entire aorta (from the ascending aorta and aortic arch to the thoracic aorta) from each group was isolated and snap-frozen in liquid nitrogen. The nuclei were isolated, following the previously published protocol. In brief, the frozen aortic tissues were cut into <5mm pieces and homogenized using a glass Dounce tissue grinder (#D8938, Sigma-Aldrich, USA). The tissues were homogenized 25 times with pestle A and 25 times with pestle B in 2ml pre-cold nuclei EZ lysis buffer. The samples were incubated on ice for 5 minutes with an additional 3ml of pre-cold EZ lysis buffer. Nuclei were centrifuged at 500 × g for 5 min at 4°C, washed with 5 ml of ice-cold EZ lysis buffer, and incubated on ice for 5 min. Following centrifugation, the nuclear pellet was washed with 5 ml of nuclei suspension buffer (NSB; 1xPBS, 0.01% BSA, and 0.1% RNase inhibitor (#2313A, Clontech, USA). The isolated nuclei were then resuspended in 2 ml of NSB, filtered through a 35 μm cell strainer (#352235, Corning Falcon, USA), and counted. A final concentration of 1,000 nuclei per µl was prepared for loading onto a 10x Genomics channel.

### Single-nucleus RNA-seq Library Preparation and Sequencing

Single-nucleus RNA-seq libraries were prepared using the Chromium Next GEM Single Cell 3ʹ Reagent Kits v3.1 on the Chromium Controller (#PN-1000121, 10x Genomics, USA). Single-nucleus suspensions were obtained from cultured cell lines, and nuclei were resuspended in PBS containing 0.04% BSA. The nuclei suspension was then loaded onto a Chromium Next GEM Chip G, and the Chromium Controller was run to generate single-cell gel beads in emulsion (GEMs) according to the manufacturer’s instructions. Within each GEM, captured nuclei were lysed, and the released RNA was barcoded through reverse transcription. Barcoded full-length cDNA was synthesized, and sequencing libraries were constructed following the manufacturer’s protocol. Library quality was assessed using a Qubit 4.0 Fluorometer and an Agilent 2100 Bioanalyzer (Agilent, USA). Sequencing was performed on an Illumina NovaSeq 6000 platform (Biomarker Technologies Corporation, Beijing, China) with a depth of at least 50,000 reads per nucleus, using 150 bp paired-end (PE150) reads.

### Data Analysis

#### Processing of Raw Data

Sequence alignment was performed using *Cell Ranger* v7.0 (10x Genomics, USA), which employs the STAR aligner. Reads were mapped to the mouse reference genome (mm10). Low-quality nuclei were excluded by retaining only those with more than 500 detected genes and a mitochondrial gene ratio below 0.20. Genes expressed in at least ten nuclei were included for downstream analysis. Gene expression was quantified using unique molecular identifiers (UMIs) assigned to each cell barcode–gene pair. After alignment, cell barcodes corresponding to valid nuclei were identified and retained using the *Cell Ranger* v7.0 pipeline.

#### scRNA-seq processing

Single-cell data were preprocessed, quality-controlled, normalized, and clustered using Seurat v4.1.3. Putative doublets were removed with *DoubletFinder* v2.0.3. QC thresholds included ≥400 detected genes per cell and ≤20% mitochondrial gene content. Subsequent normalization, variable gene selection, dimensionality reduction, and clustering followed Seurat default workflows. Batch integration across samples was performed with Harmony v1.2.3. Cell clusters were annotated using curated marker genes from literature and manual curation. Differential expression between subclusters was computed with *FindAllMarkers* (Wilcoxon test), with adj. *p*-value < 0.05 and |log_2_(Fold Change)| > 0.5.

#### MiloR

Single-nucleus data were integrated with Harmony in Seurat, factor levels were ordered and pruned, and the object was converted to *SingleCellExperiment* and initialized as a *Milo* object. A k-nearest neighbor graph (k = 20) was constructed in a 40-dimensional embedding, and overlapping neighborhoods were generated and refined using Milo (prop = 0.15, k = 20, d = 40, refinement_scheme = “graph”). Neighborhood counts were aggregated at the sample level, and a sample-level design table was aligned to the neighborhood count matrix. Differential abundance was assessed with *Milo*’s *testNhoods* using a model of the form ∼ factor, applying graph-overlap-aware FDR weighting. Visualization and QC included an abstract neighborhood graph, p-value histograms, and log_2_(Fold Change) versus −log_10_(FDR) plots, with effects mapped onto low-dimensional embeddings using *plotReducedDim* (cell-type coloring) and *plotNhoodGraphDA* with a symmetric divergent color scale. Neighborhoods were assigned to dominant cell identities via *annotateNhoods*, and results were summarized with *plotDAbeeswarm* plus boxplots referenced to zero.

#### Pseudotime trajectory analysis

*Monocle2* (v2.34.0) was used to reconstruct lineage trajectories. High–dimensional transcriptomes were embedded into a lower–dimensional manifold, enabling cells to be ordered along branching paths. Pseudotime-resolved expression dynamics were visualized with *plot_pseudotime_heatmap*.

#### Gene Set Enrichment Analysis (GSEA)

Gene Set Enrichment Analysis (GSEA) was conducted with *GSEApy*. Differential genes were ordered by decreasing log2 fold change to generate a preranked list, and Hallmark pathways (MSigDB v2024.1, h.all) were obtained via *gseapy.Msigdb()*. Enrichment was run with *gseapy.prerank* using 1,000 permutations, considering gene sets of size 1∼10,000, and reporting NES, nominal P, and FDR q for each pathway; a fixed seed was used for reproducibility.

#### pySCENIC Transcription factor analysis

We reconstructed gene regulatory networks with *pySCENIC*. Loom-formatted expression matrices and a curated catalog of human transcription factors served as inputs for GRNBoost2 to detect TF-target co-expression modules. Networks were pruned by motif enrichment in the ctx step using the TF-motif ranking database and motif annotation file, with dropout masking and parallelization enabled. Regulon activities were then scored per cell with AUCell to produce a regulon-by-cell activity matrix.

#### hdWGCNA

High-dimensional WGCNA was applied to GFP-positive cells from our data. Starting from a Seurat object, metacells were constructed by pooling cells within epithelial subtypes (k = 25; assay = SCT; max_shared = 20; min_cells = 300; reduction = umap; slot = data). We built coexpression networks using *ConstructNetwork* with a soft-threshold power of 14, and functionally annotated the resulting modules via *EnrichR*.

#### GeneTrajectory

We applied *GeneTrajectory* to all GFP-positive cells to infer gene-level dynamics without constructing cell pseudotime. Briefly, single-cell transcriptomes were preprocessed in *Scanpy*, and a cell-cell kNN graph was built to compute graph shortest-path distances.

Pairwise gene–gene distances were then obtained as optimal transport (Wasserstein) distances between gene expression distributions over the cell graph. Using these distances, genes were embedded with Diffusion Maps, and sequential gene trajectories were extracted by iteratively identifying termini and performing random-walk-based selection on the gene graph. For each trajectory, genes were ordered to yield a *pseudotemporal* ranking, where larger *pseudoorder* values indicate earlier activation. We then focused on two transitions, endothelial to Adipo-like and endothelial to SMC-like, and reported the ordered gene programs activated along each path. Visualization of trajectory bins across the embedding used smoothed expression values and gene-bin scoring to map early-to-late gene activity over the cell manifold. All analysis code followed the official *GeneTrajectory* tutorial and documentation.

#### CellOracle

*CellOracle* is a GRN-based framework for virtual gene perturbation that integrates single-cell transcriptomes with regulatory networks and, optionally, scATAC-seq co-accessibility. By coupling reconstructed GRNs with single-cell data, it models how perturbing a gene alters downstream programs and cell identity. In this work, we applied *CellOracle* to our scRNA-seq dataset using the mouse GRN distributed in the official tutorial and default settings. For in silico knockouts, the target TF’s expression was clamped to zero and its regulatory influence was iteratively propagated through the network up to five layers to approximate downstream transcriptional effects, after which *CellOracle* predicted the resulting shifts in cell-state trajectories. All analysis code followed the official tutorial and documentation.

## Statistical Analysis

All statistical analyses and visualizations were performed in R (version 4.1.3). Pearson’s correlation coefficient assessed associations between continuous variables. For quantitative comparisons, two-tailed unpaired Student’s *t* test or one-way ANOVA with Tukey’s multiple-comparisons test were used to compare subgroup means. A *p*-value < 0.05 was considered statistically significant.

## Results

### Single-Cell Lineage Tracing of Endothelial Cells in Atherosclerosis

To trace the lineage of endothelial cells (ECs) during atherosclerosis development, we generated EC-specific lineage-tracing mice (*Cdh5-CreER^T^*^2^*/Apoe^−/−^/Rosa26-mTmG*; hereafter referred to as EC reporter) ^25,26^. Tamoxifen was administered to induce *Cre* recombinase activity, followed by 12 weeks of high-fat diet (HFD) feeding to promote atherosclerotic plaque formation. Given that female sex hormones have been reported to confer vascular protection ^27^, an ovariectomized female group was included for comparison. In total, five experimental groups were analyzed: HFD-fed male EC reporter (EM), HFD-fed female EC reporter (EFe), ovariectomized HFD-fed female EC reporter (OEEFe), and chow diet-fed *Rosa26-mTmG* male (mM) and female (mFe) controls. For the main experimental groups (EM and EFe), single-nucleus RNA sequencing (snRNA-seq) was performed in biological triplicates, with each replicate consisting of pooled aortae from at least three mice. For the OEEFe, mM, and mFe groups, pooled aortae from a minimum of three mice were subjected to a single snRNA-seq experiment per group (**Figure 1A**).

**Figure 1.**
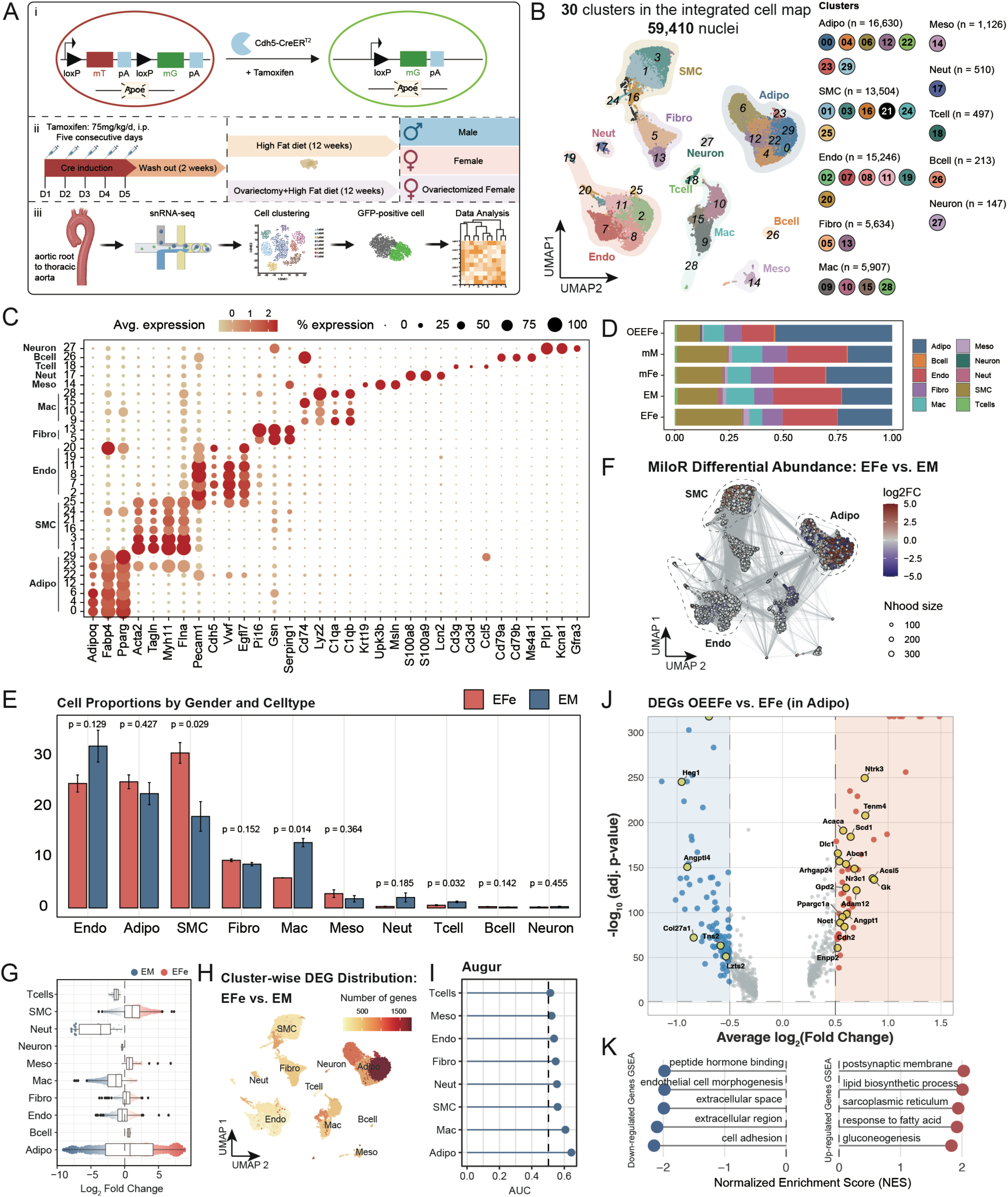
Single-cell lineage tracing of endothelial cells in atherosclerosis. **A.** Schematic illustration of the generation of EC-specific reporter mice, the atherosclerosis model, and the single-cell lineage-tracing strategy. **B.** Uniform manifold approximation and projection (UMAP) visualization of all nuclei integrated from all samples. **C.** Dot plot of canonical markers used for cell annotation. Dot size represents the fraction of cells expressing each gene; color gradient indicates average expression. **D.** Comparison of cell-type composition of indicated samples. **E.** Comparison of the proportions of different cell types between the EFe and EM groups. Within each cell type, pairwise comparisons were performed between groups. Two-sided unpaired Student’s t-test was used for statistical analysis; error bars represent mean ± SD. Exact p values are displayed above each comparison on the plot. **F.** The plot of differential abundance between EFe and EM group by *MiloR*. **G.** The violin plots show neighborhood-level log fold changes across major cell types between EFe and EM. **H.** Differential expression landscape between sexes. UMAP highlighting clusters with significant DE (adjusted p<0.05) and per-cluster DEG counts. **I.** Augur perturbation analysis indicates differential perturbation in various cell types using area under the classifier curve (AUC), the dash line equals 0.5. **J.** The volcano plot for comparing differential expressed genes of adipocytes between OEEFe and EFe groups. Differential expressions were calculated using Wilcoxon rank-sum test. P values were adjusted for multiple testing using the Benjamini-Hochberg procedure to control the false discovery rate (FDR) and are reported as adjusted p values (adj. p-value). Significance was defined as adj. p < 0.05 and |log2FC| ≥ 0.5. **K.** GSEA of adipocyte DEGs showing enrichment of lipid biosynthesis, cholesterol handling, and related pathways.

We profiled a total of 59,410 single nuclei and 30 clusters from all groups. Using established marker genes ^28^, we identified 16,630 adipocytes (*Adipoq*⁺, *Fabp4*⁺, *Pparg*⁺), 15,246 endothelial cells (*Pecam1*⁺, *Cdh5*⁺, *Vwf*⁺, *Egfl7*⁺), 13,504 smooth muscle cells (*Acta2*⁺, *Tagln*⁺, *Myh11*⁺, *Flna*⁺), 5,907 macrophages (*Cd74*⁺, *Lyz2*⁺, *C1qa*⁺, *C1qb*⁺), 5,634 fibroblasts (*Pi16*⁺, *Gsn*⁺, *Serping1*⁺), 1,126 mesothelial cells (*Msln*⁺, *Upk3b*⁺, *Krt19*⁺), 510 neutrophils (*S100a8*⁺, *S100a9*⁺, *Lcn2*⁺), 497 T lymphocytes (*Cd3g*⁺, *Cd3d*⁺, *Ccl5*⁺), 213 B lymphocytes (*Cd79a*⁺, *Cd79b*⁺, *Ms4a1*⁺), and 147 neurons (*Plp1*⁺, *Kcna1*⁺, *Gfra3*⁺) (**Figure 1B-1C; Supplemental Dataset S1**). Comparative analysis of cell-type composition revealed sex-specific differences: males exhibited significantly higher proportions of macrophages (p=0.014) and T lymphocytes (p=0.032), consistent with prior clinical observations of sex differences in atherosclerotic plaques (**Figure 1D-1E**) ^10^. While females showed a greater abundance of SMCs (p=0.029).

To further dissect these differences, we applied sex-specific differential abundance testing across all cell types using *miloR* ^29–31^, which revealed widespread sex-dependent alterations across multiple cellular neighborhoods (**Figure 1F-1G**). Differential expression analysis (adjusted p<0.05) revealed extensive transcriptional perturbations between sexes, with adipocytes being the most transcriptionally affected population (**Figure 1H**). This finding was further validated using the *Augur* algorithm. This machine-learning framework quantifies cell-type-specific perturbations, which again identified adipocyte-like cells as the most strongly perturbed cell population (**Figure 1I**) ^32^.

Given that we observed a marked increase in the proportion of adipocytes (Adipo) in the OEEFe group versus the EFe group (from 25.10% to 53.76%) (**Figure 1D**), suggesting a critical role of adipocytes in the progression of atherosclerosis in menopausal females after the loss of female sex hormones ^33,34^. Comparison of adipocytes from ovariectomized (OEEFe) and intact female (EFe) mice revealed more than 650 differentially expressed genes (DEGs) (**Figure 1J-1K, Supplemental Dataset S2**). Upregulated genes were predominantly enriched in pathways related to lipid biosynthesis, cholesterol efflux, and transcriptional reprogramming, including *Pparg*, *Srebf1*, *Ppargc1a*, *Acly*, *Acaca*, *Fasn*, and *Acsl5*, indicating an active lipogenic and *de novo* fatty acid synthesis state ^35,36^. In contrast, downregulated genes such as *Angptl4*, *Heg1*, *Col27a1*, *Tns2*, and *Lzts2* indicated attenuation of matrix/adhesion and vascular-protective signals that typically restrain lipid flux and tissue remodeling. These findings suggest a shift toward increased lipid anabolism and decreased catabolism following the loss of female hormones, consistent with clinical observations of accelerated plaque growth after menopause ^37^.

### Sex-difference of EC-derived Subpopulation in Atherosclerosis

To investigate sex-biased endothelial cell (EC) trajectories in atherosclerosis, we analyzed single-nucleus data by selecting cells annotated as ECs or EC-derived populations. We subsequently re-clustered a total of 15,246 ECs and identified nine distinct EC subclusters (**Figure 2A**). To annotate these subtypes, a marker-gene-based scoring system was developed using previously reported EC markers for scRNA-seq ^38^. We defined three major EC subtypes: arterial ECs (Art_EC), marked by *Efnb2*, *Sox17*, *Bmx*, *Sema3g*, *Hey1*, *Ltbp4*, *Fbln5*, *Gja5*, *Gja4*; capillary ECs (Cap_EC) defined by *Car4*, *Prx*, *Rgcc*, *Sparc*, *Sgk1*; and lymphatic ECs (Lym_EC) characterized by *Prox1*, *Lyve1*, *Flt4*, and *Pdpn*, whereas venous ECs (Ven_EC) were not found in our data by *Nr2f2*, *Vcam1*, *Ackr1*, *Selp* (**Figure 2B-2C; Supplemental Dataset S3).** Notably, two clusters were annotated to capillary ECs (Cap_EC) and lymphatic ECs (Lym_EC), whereas the remaining seven EC subclusters were annotated based on their dominant signature genes as Art_EC-*Fmo2*^+^, Art_EC-*Col8a1*^+^, Art_EC-*Ly6a*^+^, Art_EC-*Sox17*^+^, Art_EC-*Myh11*^+^, Art_EC-*Mki67*^+^, and Art_EC-*Myh6*^+^ (**Figure 2C-2D; Supplementary Figure S1A**). Art_EC-Ly6a⁺ is enriched in vascular progenitors and activated ECs, contributing to the formation of new blood vessels in response to injury, which is a key process in atherosclerosis (**Figure 2E-i**) ^39–41^; Art_EC-*Fmo2*⁺ is associated with redox balancing, NO sensing and injury repair (**Figure 2E-ii**) ^42^; Art_EC-*Col8a1*⁺ cells were marked by *Col8a1*, *Fbln5*, and *Eln*, showing a role in ECM deposition and plaque stability (**Figure 2E-iii**) ^43,44^; Art_EC-*Sox17*⁺ cells maintain the EC identity as a key transcription factor in EC lineage commitment (**Figure 2E-iv**) ^45,46^; Art_EC-*Myh11*⁺ cells express SMC-specific myosin heavy chain, representing an EC-to-SMC transition subpopulation that may play a crucial role in fibrous cap development and stability (**Figure 2E-v**) ^47^; Art_EC-*Mki67*⁺ cells representing a proliferative reservoir of arterial endothelium (**Figure 2E-vi**) ^48^; Art_EC-*Myh6*⁺ cells distribute near the aortic root, potentially adapted to the compliance demands near the aortic outflow tract (**Figure 2E-vii**) ^49^.

**Figure 2.**
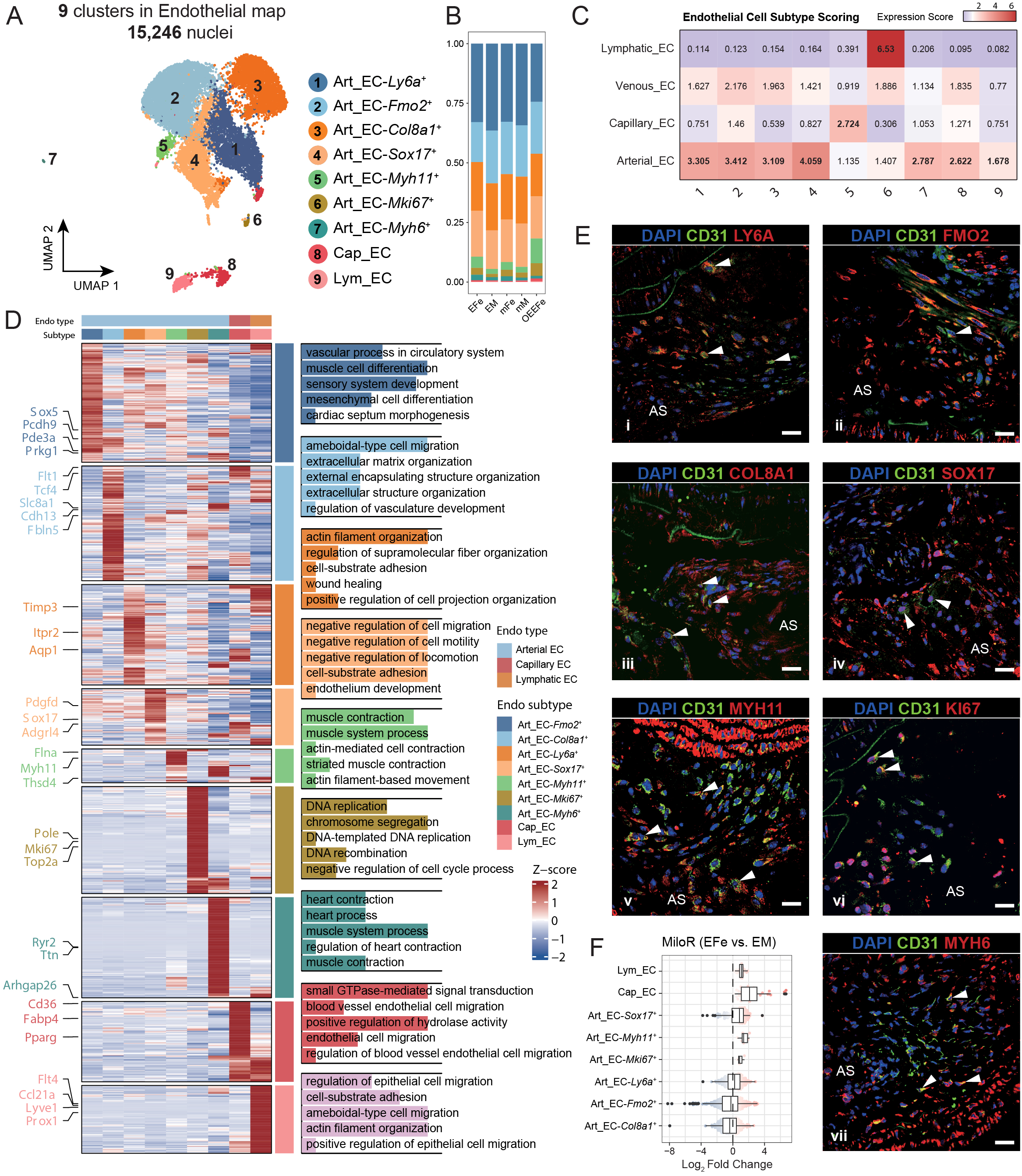
Sex-difference of EC-derived subpopulation in atherosclerosis. **A.** UMAP of all cells annotated as EC by canonical cell markers (*Pecam1*, *Cdh5*, *Vwf*, *Egfl7*), a total of 15,246 nuclei was re-clustered into 9 subclusters. **B.** Group-wise composition of all EC subclusters. **C.** Marker-based subtype scoring using published EC signatures, confirming arterial, capillary, and lymphatic identities; venous signatures were not detected. **D.** Heatmap of representative genes and enriched GO terms delineating biological programs for each EC state. **E.** Immunofluorescence validation of the existence of different EC subtypes in the atherosclerotic plaque, scale bar indicates 20μm. **F.** Differential abundance between EFe and EM by MiloR in all EC subclusters. **G.** Augur perturbation analysis indicates differential perturbation in different EC subtypes using area under the classifier curve (AUC), the dash line equals 0.5.

To study the sex difference of these EC subtypes in atherosclerosis, the *MiloR* algorithm was used to analyze the differential abundance (DA) of each subcluster ^50^, and the results indicated that more Cap_EC, Lym_EC, Art_EC-*Sox17*^+^, Art_EC-*Myh11*⁺, Art_EC-*Mki67*^+^, and Art_EC-*Ly6a*^+^ are enriched in female atherosclerotic aorta, whereas Art_EC-*Col8a1*⁺ and Art_EC-*Fmo2*⁺ are more abundant in males (**Figure 2F**). To quantitatively determine which subclusters differ between males and females, we applied the *Augur* algorithm ^32^. The results indicated that the Art_EC-*Myh11⁺* and Art_EC-*Ly6a⁺* subclusters were preferentially enriched in females (**Figure 2G; Supplementary Figure S1B-S1K**).

To characterize the functional features of each EC subcluster, particularly Art_EC-*Myh11*⁺ and Art_EC-*Ly6a*⁺, we performed differential gene expression (DEG) and gene ontology (GO) analyses. Our data revealed that female Art_EC-*Myh11*⁺ cells exhibited downregulation of fatty acid metabolic processes and cell migration compared with males (**Supplemental Figure S2A**), whereas Art_EC-*Ly6a*⁺ cells showed reduced angiogenesis and enhanced cell adhesion (**Supplementary Figure S2B**). Functional analyses of other EC subtypes are also presented (**Supplementary Figure S2C-S2D and S3A-S3E**). Collectively, these findings suggest sex-specific EC transdifferentiation toward distinct cell lineages, which may contribute to the differential characteristics of atherosclerotic plaques.

### Sex Differences in Gene Expression of EC-derived Cells in Atherosclerosis

Similar to SMCs, which exhibit high plasticity and undergo phenotypic switching under physiological or pathological conditions ^13,51^, ECs also display considerable, though comparatively limited, plasticity ^19,52,53^. However, sex-dependent differences in the role of ECs during atherosclerosis development remain poorly understood ^54^.

Here, we identified 6,113 GFP⁺ cells, accounting for 11.5% of the entire dataset (**Figure 3A**). In EFe and EM groups, 4,796 cells (78.39%) were annotated as ECs, indicating that 21.61% of GFP⁺ cells no longer expressed EC markers, suggesting that these cells had undergone transdifferentiation into other cell types (**Figure 3B**). Our data showed that 7.65% of GFP⁺ cells transdifferentiated into adipocyte-like cells (468 cells), 6.18% into SMC-like cells (378 cells), 3.11% into macrophage-like cells (190 cells), and 2.93% into fibroblast-like cells (179 cells) (**Figure 3B; Supplementary Dataset S4**). Notably, the proportion of transdifferentiated ECs was higher in females (730/2,940, 24.83%) compared to males (592/3,178, 18.63%) (**Figure 3C**). Specifically, 9.93% of EC-derived GFP⁺ cells in females (292/2,940) acquired adipocyte-like features, compared to 5.54% in males (176/3,178). In contrast, approximately 7.96% of ECs transdifferentiated into SMC-like cells in females, whereas 4.65% underwent this transition in males (**Figure 3C**).

**Figure 3.**
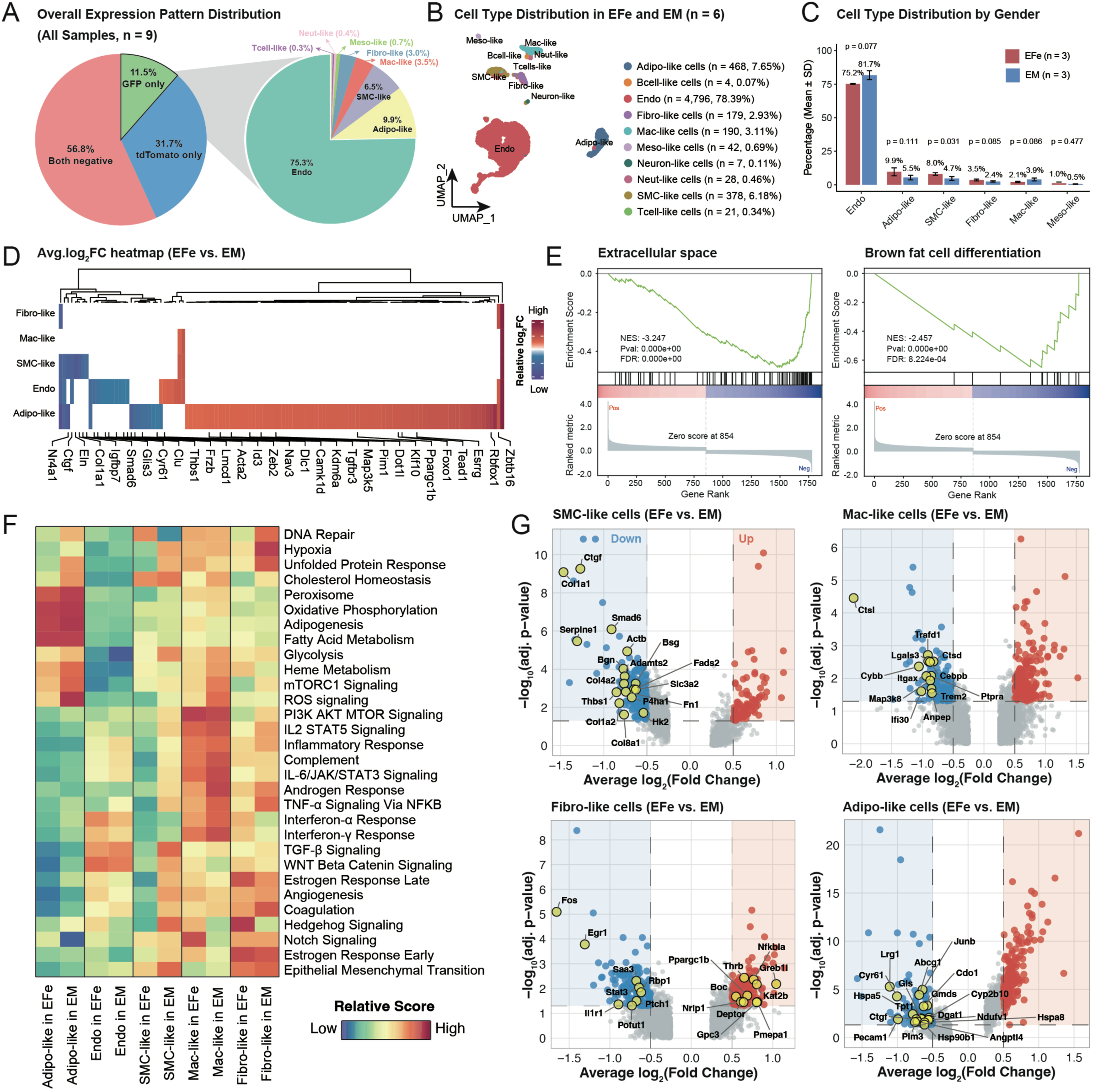
Sex differences in gene expression of EC-derived cells in atherosclerosis. **A.** The pie chart shows that GFP⁺ cells account for 11.5% of all cells, and these GFP⁺ nuclei can be further classified into various EC-derived transdifferentiated cell types. **B.** UMAP of GFP⁺ cells colored by lineage identity (left) and cell proportion of each cell subtypes derived from GFP⁺ cells. **C.** Comparison of different cell subtype proportions derived from GFP⁺ cells stratified by sex (EFe vs. EM), pairwise comparisons were performed between groups. Two-sided unpaired Student’s t-test was used, and exact p values are displayed above each comparison on the plot. **D.** Heatmap of average log2 fold change genes (EFe vs EM) for each EC-derived lineage with representative genes labeled. **E.** GSEA enrichment plots for adipocyte-like cells comparing females and males, displaying running enrichment scores and ranked gene lists for two example pathways. **F.** Heatmap of GSVA hallmark pathway scores across EC-derived lineages by sex. **G.** Volcano plots of differential expression (EFe vs EM) for SMC-like, macrophage-like, fibroblast-like, and adipocyte-like cells with selected genes annotated. Differential expressions were calculated using Wilcoxon rank-sum test. P values were adjusted for multiple testing using the Benjamini-Hochberg procedure to control the false discovery rate (FDR) and are reported as adjusted p values (adj. p-value). Significance was defined as adj. p < 0.05 and |log2FC| ≥ 0.5.

Next, to investigate sex differences in EC-derived cells, we analyzed the gene expression profiles of each cell subtype. The results revealed minimal differences between fibroblast-like and macrophage-like cells from males and females, whereas SMC-like cells and ECs exhibited moderate sex-dependent variation. Notably, adipocyte-like cells displayed the most distinct gene expression patterns between the two sexes (**Figure 3D; Supplementary Dataset S5**). GSEA analysis of adipose-like cells revealed significantly reduced enrichment of pathways related to the extracellular space (NES = −3.247, FDR<0.05) and brown fat cell (BAT) differentiation (NES = −2.457, FDR<0.05) in females compared to males (**Figure 3E**).

To further dissect sex differences, we performed gene set variation analysis (GSVA) across EC-derived lineages ^55^. The SMC-like cells from males exhibited higher activity in glycolysis, TGF-β signaling, and Hedgehog signaling pathways (**Figure 3F; Supplementary Dataset S6**), suggesting that these cells adopt a synthetic phenotype characterized by active proliferation and dedifferentiation ^56,57^. The macrophage-like cells from males showed broad activation of the TNF-α, IL-6/JAK/STAT3, and interferon-α and -γ pathways, indicating that EC-derived macrophage-like cells adopt a proinflammatory, M1-polarized state ^58^. In fibroblast-like cells, females exhibit higher activity in migration-associated Hedgehog signaling and estrogen response, while responding less to inflammation-associated pathways. Interestingly, adipocyte-like cells from females exhibit lower activity in response to hypoxia, ROS signaling, and, most notably, mTORC1 signaling, suggesting that these cells are relatively more inert than their male counterparts, consistent with the less atherosclerotic plaque formation observed in premenopausal females **(Figure 3F**). To corroborate these patterns, we identified differentially expressed genes between females and males across cell types and annotated genes that contribute to the corresponding pathways **(Figure 3G, Supplementary Dataset S7-S10**).

### Sex-biased Gene Regulatory Network in EC-derived Cells during Atherosclerosis

To investigate co-expression networks underlying EC differentiation into other cell types, we applied hierarchical and dynamic weighted gene co-expression network analysis (*hdWGCNA*) to GFP⁺ cells from our dataset ^59^. This analysis identified nine distinct gene co-expression modules, designated M1-M8 (**Figure 4A, Supplemental Dataset 11**). Examination of module distribution across groups revealed distinct sex-specific patterns: Module 2 (M2) was predominantly expressed in control females (mFe), Module 5 (M5) was enriched in intact atherosclerotic females (EFe), and Modules 6 (M6) and 7 (M7) were preferentially represented in males (EM) (**Figure 4B**). Next, we performed a functional enrichment analysis of the hub genes and identified distinct roles for the individual modules. Module 2 was enriched in pathways related to lipid and fatty acid metabolism, predominantly in the non-atherosclerotic group, suggesting that Module 2 plays a critical role in maintaining baseline fatty acid metabolic programs essential for vascular homeostasis. Module 5, selectively enriched in female atherosclerotic tissue (EFe), involved cellular responses to laminar shear stress and adhesion, representing a mechanical defense network that may contribute to delayed onset and enhanced plaque stability in females, consistent with previous reports ^60–62^. Modules 6 and 7, activated mainly in male atherosclerotic tissue, were associated with extracellular matrix organization (M6) and regulation of SMC proliferation and migration (M7), reflecting the pronounced SMC phenotypic switching and plaque formation in males and highlighting the importance of EC-SMC crosstalk (**Figure 4C, Supplemental Dataset 12**) ^13^. To elucidate the internal relationships among these modules, hub genes network visualization was performed.

**Figure 4.**
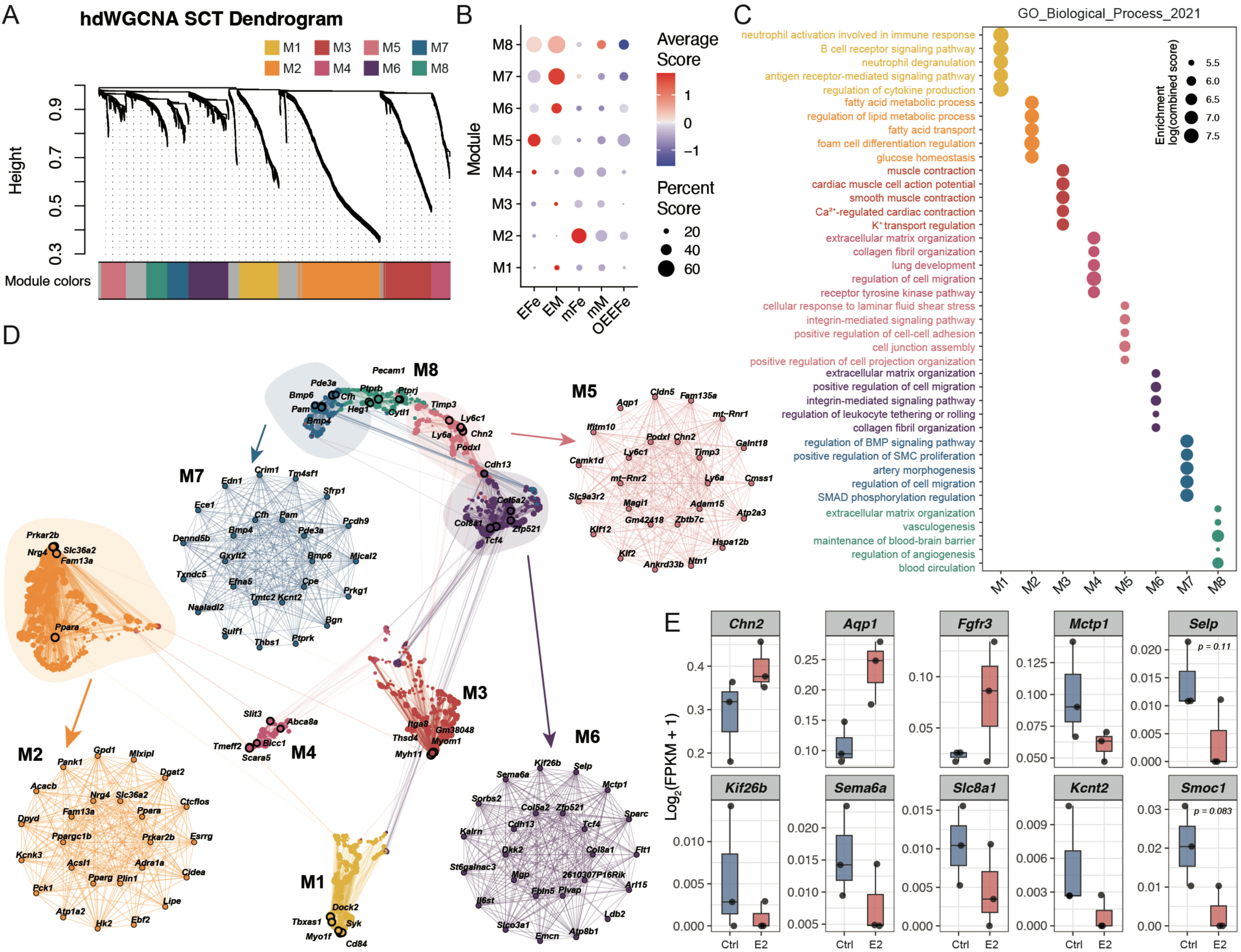
Sex-biased gene co-expression modules and hub networks in EC-derived GFP+ cells. **A.** hdWGCNA dendrogram of GFP⁺ cells indicate nine co-expression modules (M1-M8) colored by module assignment. **B.** Dot plot summarizing module expression across groups. Dot size indicates the score of cells expressing module genes; color denotes average expression. **C.** GO Biological Process (GO-BP) enrichment for hub genes of each module. Dot size represents enrichment significance, color shows enrichment score. **D.** Network visualization of module-specific hub genes. Nodes represent genes; edges indicate co-expression connectivity. Module communities are outlined and labeled (M2, M5, M6, M7). **E.** Boxplots of selected genes measured by NextGen high throughput RNA-seq using Telo-HAEC cell line under vehicle control (Ctrl) or estradiol (E2) treatment.

To identify the key nodes driving sex-specific network differences, we examined and visualized the highest-connectivity hub genes within each co-expression module (**Figure 4D, Supplemental Dataset 12**). In M5, genes such as *Klf2* and *Cldn5* have been reported to be upregulated by estrogen and are known to alleviate endothelial inflammation and oxidative stress ^63,64^. In M6, genes including *Col8a1*, *Fbln5*, and *Mgp* are associated with inflammatory Endo-MT, vascular oxidative environment, and calcification ^65–67^. In M7, genes such as *Bmp6* and *Thbs1* participate in vascular calcification, plaque rupture, and endothelial dysfunction ^68,69^, forming the molecular basis for plaque progression, instability, and calcification in males. Interestingly, module M2, specific to healthy females, is enriched for known atheroprotective genes, including *Pparg*, *Ppara*, and *Ppargc1b*. These genes confer protection by promoting fatty acid oxidation, regulating cholesterol metabolism, suppressing vascular inflammation, and maintaining vascular energy homeostasis ^70,71^.

To investigate whether the regulatory network is influenced by female hormones, we treated cultured human endothelial cells (Telo-HAEC) with estradiol and performed RNA sequencing. Integrated analysis of the snRNA-seq and RNA-seq data revealed that genes within the regulatory network are modulated by estrogen, validating the findings of sex-specific EC regulatory network in atherosclerosis (**Figure 4E**).

### EC Transdifferentiation Trajectories in Atherosclerosis

To study the dynamics of EC differentiation in atherosclerosis, pseudotime analysis was performed using Monocle 2 ^72^, revealing three distinct cell states and two differentiation trajectories: ECs to adipocyte-like cells and ECs to SMC-like cells (**Figure 5A**). We analyzed the cellular composition of each state and found that State 1 consisted of over 98.26% ECs, State 3 comprised more than 94.65% adipocyte-like cells, whereas State 2 contained a considerable proportion of SMC-like cells (44.98%), macrophage-like cells (20.61%), and fibroblast-like cells (19.25%) (**Figure 5B, Supplemental Dataset 13**).

**Figure 5.**
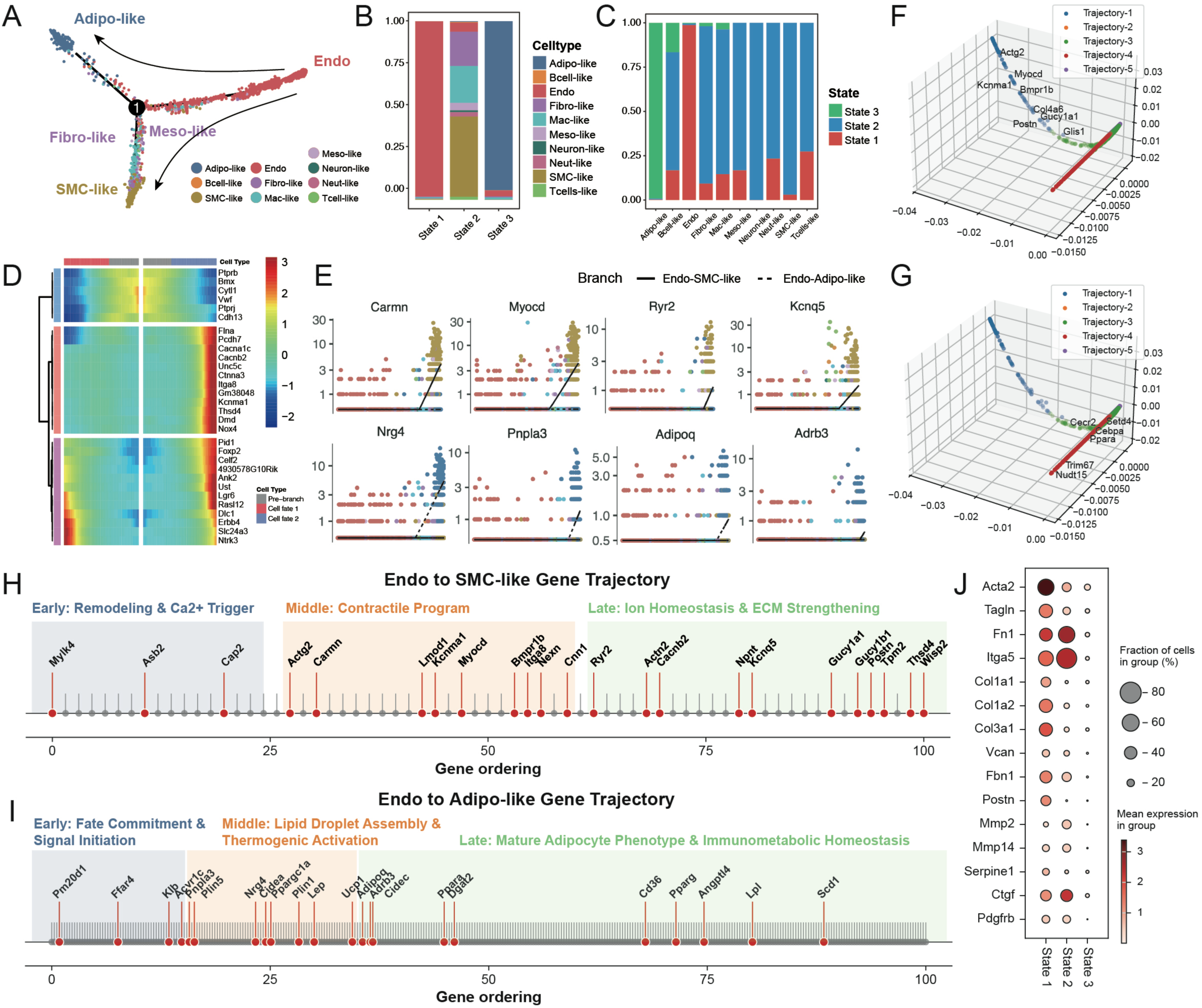
EC transdifferentiation trajectories. **A.** Monocle 2 pseudotime tree of EC and EC-derived cells showing three states and two branches, two arrows indicate different transdifferentiation preferences. **B.** Stacked bar plot of cell-type composition within each Monocle state. **C.** Proportional distribution of major EC-derived lineages in different cell states. **D.** Heatmap of pseudotime-ordered genes across states from Monocle 2. **E.** Example genes varying along the two branches, with expression plotted against pseudotime for EC→SMC-like (solid line) and EC→Adipo-like (dashed line) trajectories. **F-G.** Three-dimensional Diffusion Map embeddings of genes from GeneTrajectory, indicating differential transdifferentiation trajectories including EC to SMC-like (F) and EC to Adipo-like (G). **H.** Ordered genes along the EC to SMC-like trajectory annotated by module membership (early, middle, late). **I.** Ordered genes along with the EC to Adipo-like trajectory annotated by module membership (early, middle, late). **J.** Dot plot of canonical markers related to EndoMT programs across EC, SMC-like, and Adipo-like groups; dot size indicates fraction of expressing cells and color indicates mean expression.

Interestingly, very few fibroblast-like cells (19/5176 cells) were observed in the branch of State 3, which is predominantly composed of adipocyte-like cells, suggesting that the transdifferentiation from ECs to adipocyte-like cells is more direct compared to that from ECs to SMC-like cells (**Figure 5C**). As reported in previous studies, the transdifferentiation from ECs to SMC-like cells involves a clear endothelial-to-mesenchymal transition (EndoMT) ^19^, with fibroblast-like cells (also referred to as mesenchymal-like cells) representing an obligatory intermediate stage in the differentiation toward SMC-like cells in State 2. In contrast, the transdifferentiation of ECs into adipocyte-like cells may not necessarily undergo this intermediate step. Using Monocle 2, we furtherly characterized the differential gene expression across the various states (**Figure 5D**) and identified signature genes specifically associated with EC-to-SMC-like cell differentiation (including *Carmn, Myocd, Ryr2, Kcnq5*) and EC-to-adipocyte-like cell differentiation (including *Nrg4, Pnpla3, Adipoq, Adrb3*) (**Figure 5E**).

To further identify the key genes governing lineage commitment in the EC to SMC-like or EC to adipocyte-like cell transdifferentiation, we applied the *GeneTrajectory* algorithm, which decouples distinct gene expression programs by calculating the optimal transport distances of gene distributions across the cellular graph ^73^. The analysis revealed two major trajectories along the transdifferentiation timeframe, consistent with the results obtained from Monocle 2 (**Figure 5F-5G**). Along the SMC trajectory, we identified three gene modules: Module 1 (*Mylk4, Asb2, Cap2*) regulates cytoskeletal remodeling and calcium sensing; Module 2 (*Myocd, Carmn, Actg2, Lmod1, Kcnma1*) modulates the contractile function of SMC-like cells; and Module 3 (*Ryr2, Cacnb2, Kcnq5, Postn, Npnt*) controls ion channel activity and reinforces the extracellular matrix (ECM). These results delineate clear, stage-wise key events in the EC-to-SMC-like cell transdifferentiation process (**Figure 5H; Supplementary Dataset S14**). Along the adipocyte-like trajectory, we identified three gene modules: Module A (*Pm20d1, Ffar4, Klb, Acvr1c*) initiates the loss of endothelial identity, activates KLF2/NRF2 signaling, and suppresses MAPK/MEK-mediated pathways; Module B (*Pnpla3, Plin5, Nrg4, Ppargc1a, Ucp1*) promotes lipid droplet formation and mitochondrial biogenesis; and Module C (*Adipoq, Adrb3, Cidec, Ppara, Cd36, Pparg, Angptl4, Lpl, Scd1*) establishes a mature adipokine secretion and cholesterol efflux program (**Figure 5I; Supplementary Dataset S15**). Interestingly, our data revealed no detectable expression of genes associated with the EndoMT process, consistent with our previous hypothesis that EC-to-adipocyte-like cell transdifferentiation may bypass EndoMT, which challenges the conventional understanding.

To validate this conclusion, we examined the expression of widely reported EndoMT markers in Endo (State 1), SMC-like (State 2), and Adipo-like (State 3) cells. These included key EndoMT-related genes involved in cytoskeletal organization, collagen and matrix remodeling, and EndoMT processes (**Figure 5J**), as well as markers associated with myofibroblast and myogenic differentiation, extracellular matrix, and cell adhesion (**Supplementary Figure S4A-S4C**). These data demonstrated that the transdifferentiation from EC to adipocyte-like cells skips EndoMT, at least in the resolution of our snRNA-seq data.

### Sex-Specific Transcription Factor Governs EC Transdifferentiation in Atherosclerosis

Next, to uncover the key transcription factors (TFs) governing EC-to-adipo-like or EC-to-SMC-like cell transdifferentiation, *pySCENIC* was applied to infer gene regulatory networks from EC-derived GFP-positive cells in our model (**Figure 6A**) ^74^. Principal component analysis (PCA) of TF activation AUC scores in each cell type revealed both shared and distinct regulatory patterns, reflecting a common endothelial origin with diverging cell fates (**Figure 6B**). In adipocyte-like cells, in addition to the canonical adipogenic triad (*Pparg*, *Ppara*, *Srebf1*), the top-ranked TFs included *Esrra* and *Esrrg*, suggesting that EC-derived adipocyte-like cells are responsive to female hormones. In SMC-like cells, transcription factors including *Tead3* and *Sox9* are activated. These factors are associated with YAP/TAZ signaling and promote EndoMT that is triggered by disturbed flow (**Figure 6C**) ^75^.

**Figure 6.**
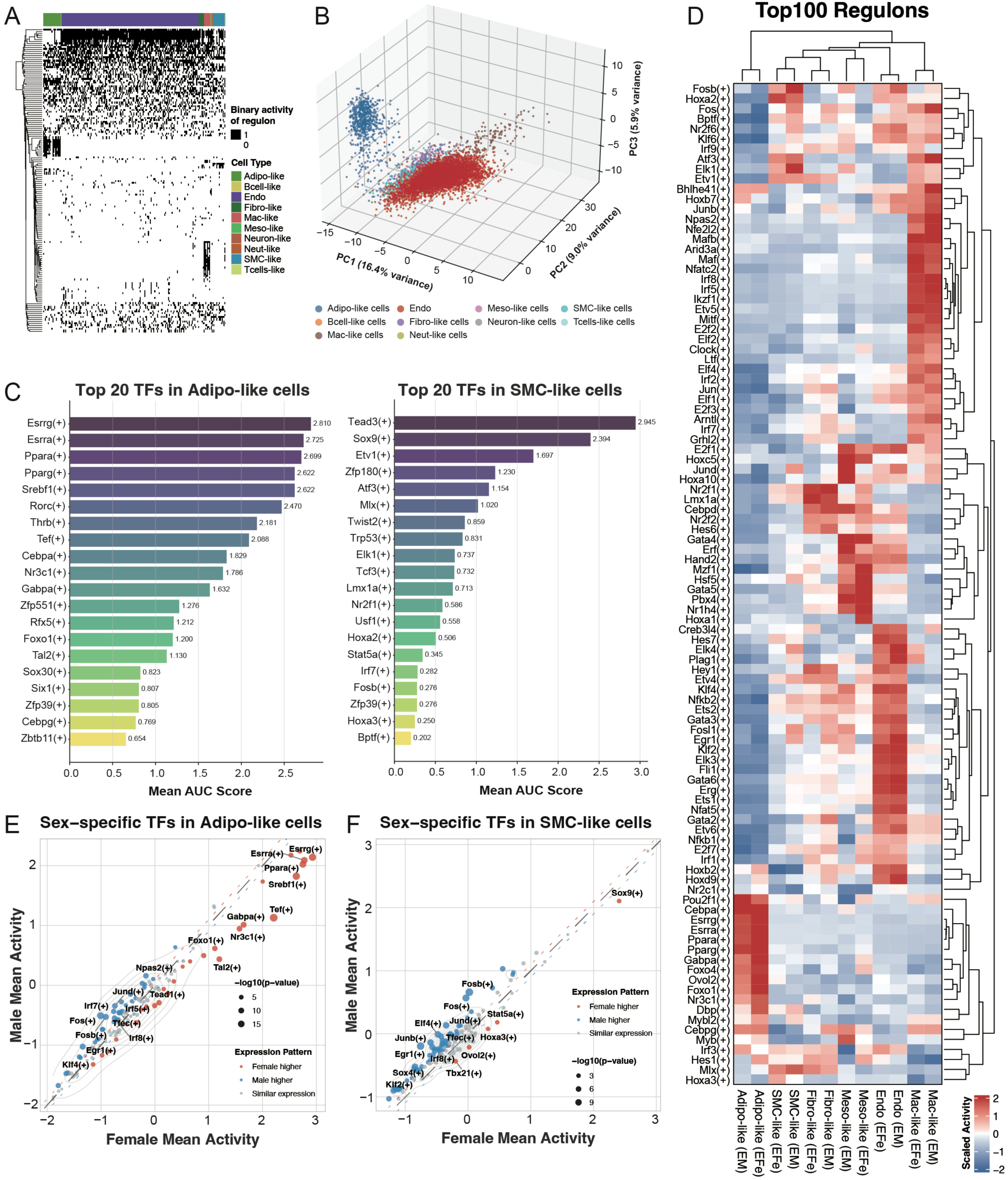
Transcription factor regulons underlying EC-derived lineages and sex-stratified activity. **A.** Binary regulon activity matrix (AUCell) across EC-derived GFP+ cells, grouped by cell type. **B.** PCA of regulon activity scores showing separation of EC, adipocyte-like, and SMC-like populations. **C.** Bar charts of the top transcription factors ranked by mean AUCell score within adipocyte-like cells (left) and SMC-like cells (right). **D.** Heatmap of the top 100 regulons across EC-derived cell types and groups, clustered by both regulons and samples. **E.** Scatter plot comparing female versus male mean regulon activity in adipocyte-like cells; point size encodes significance, and color denotes relative bias. **F.** Scatter plot comparing female versus male mean regulon activity in SMC-like cells; point size encodes significance, and color denotes relative bias. P values were computed per TF within each cell type using a two-sided unpaired Student’s t-test, Benjamini-Hochberg correction was then applied across TFs within each cell type to obtain adjusted p values (adj. p-value).

To investigate the sex-biased TFs in EC transdifferentiation, we compared the top 100 TF regulons along each differentiation trajectory (**Figure 6D, Supplementary Dataset 16**). In adipocyte-like cells, females showed higher activation of the metabolic-steroid receptor module, including *Esrra*, *Esrrg*, *Ppara*, *Srebf1*, *Gabpa*, *Foxo1*, and *Nr3c1*, whereas males preferentially activated the *Jun*/*Fos* and *Irf7*/*8* axes associated with stress and inflammation (**Figure 6E**). These results suggest that female adipocyte-like cells are more inclined toward metabolic homeostasis and antioxidative functions, while their male counterparts are more prone to pro-inflammatory and stress-responsive states. In SMC-like cells, males exhibited stronger activation of transcription factors associated with phenotypic switching, vascular injury responses, and inflammation, including the AP-1 family (*Fos*, *Fosb*, *Junb*, *Jund*) and *Egr1* ^76^. By contrast, female SMC-like cells preferentially expressed *Sox9*, a regulator that enhances ECM stiffness and promotes SMC senescence, suggesting that males and females may undergo phenotypic switching through distinct molecular mechanisms (**Figure 6F**) ^77^.

Next, we screened TFs that met dual criteria of both sex specificity and transdifferentiation association, yielding 49 TFs in the adipogenic branch and 45 TFs in the SMC branch, resulting in 94 non-redundant candidate regulators (**Figure 7A, Supplementary Dataset 17**). These TFs were subjected to *in silico* one-by-one knockout analysis using *CellOracle* (**Figure 7B**) ^78^. Our data indicated that virtual deletion of *Klf2* significantly redirected the differentiation trajectory toward the SMC branch, with modest expansion into the adipocyte-/macrophage-like regions, demonstrating KLF2 acts as a homeostatic “brake” on EndoMT (**Figure 7C**) ^79,80^. Furthermore, virtual deletion of *Esrrg* promotes SMC trajectories and inhibition of adipogenic differentiation (**Figure 7D**). Correlation analysis further supported these functional roles: *Esrrg* was positively correlated with *Ucp1, Ppargc1a/b*, *Cidea*, *Nrg4*, and *Plin5*, genes characteristic of brown/beige fat thermogenesis and energy dissipation (**Figure 7F, Supplementary Figure S5A-S5I**). In contrast, *Creb5* deletion inhibited SMC differentiation while promoting adipogenic transition (**Figure 7E**). Moreover, *Creb5* expression correlated with *Col1a2*, *Fn1*, and *Gata6*, genes involved in extracellular matrix remodeling and SMC lineage specification (**Figure 7F, Supplementary Figure S5A-S5I**). Considering that CREB family members respond to both shear stress and cAMP signaling ^81^, these findings suggest that CREB5 may act as a mechanosensitive regulator linking mechanical cues to smooth muscle fate commitment.

**Figure 7.**
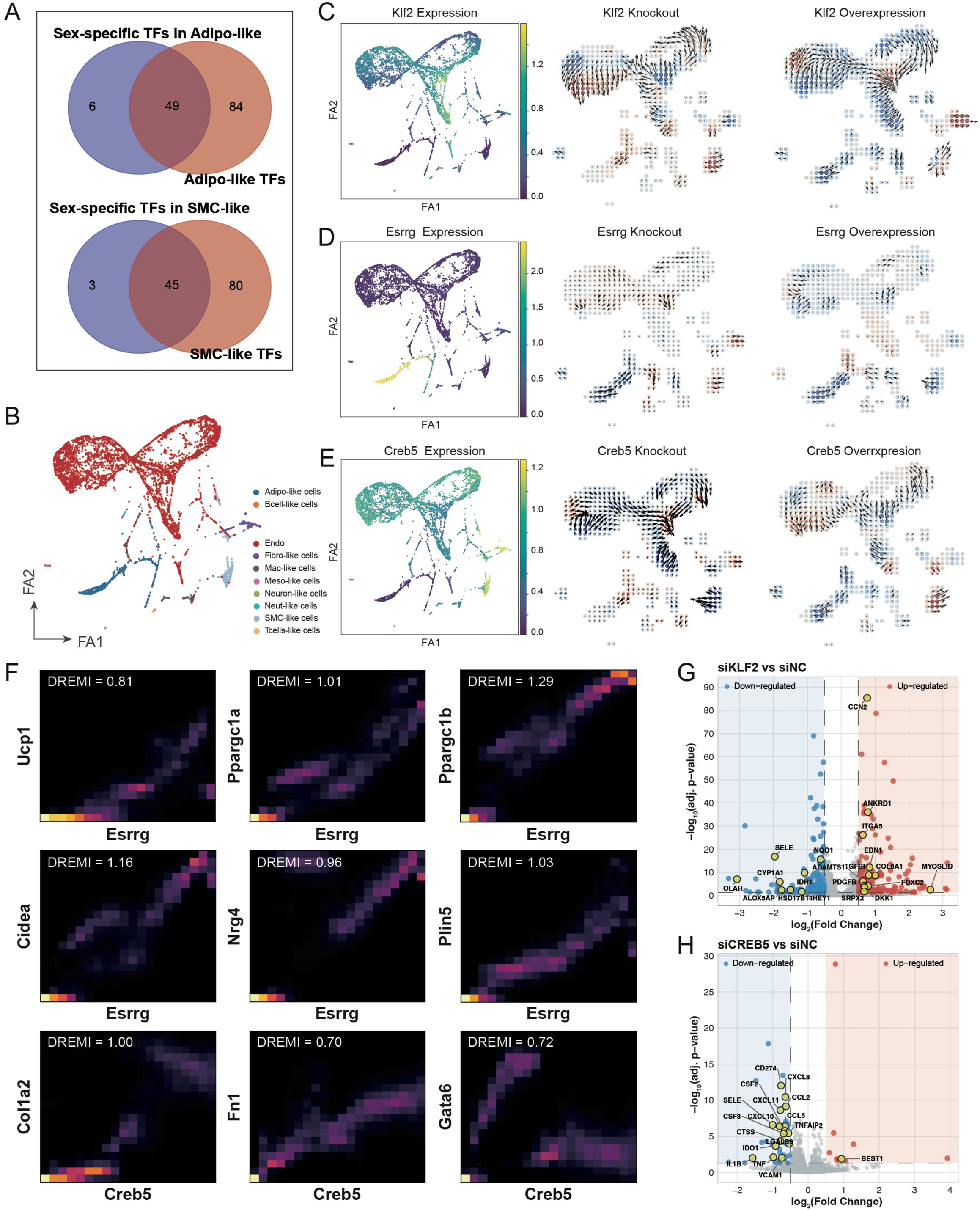
Candidate transcription factors driving EC transdifferentiation and *in silico* perturbation. **A.** Venn diagrams showing TFs meeting dual criteria of sex specificity and trajectory association for the adipocyte-like branch and the SMC-like branch, and the combined non-redundant set. **B.** Force-directed embedding of EC-derived cells colored by lineage identity used for CellOracle simulations. **C-E.** CellOracle in silico perturbations for *Klf2* (C), *Esrrg* (D), and *Creb5* (E). For each TF, left panels show baseline expression over the manifold; middle and right panels display predicted fate probability shifts under knockout or overexpression, respectively. **F.** DREMI/conditional probability plots illustrating associations between *Esrrg* or *Creb5* expression and selected target or marker genes. **G-H.** Volcano plots depict differentially expressed genes for siKLF2 versus siNC (G) and siCREB5 versus siNC (H), with representative genes labeled, based on NextGen high throughput RNA-seq with siRNA knockdown. Differential expressions were calculated using Wilcoxon rank-sum test. P values were adjusted for multiple testing using the Benjamini-Hochberg procedure to control the false discovery rate (FDR) and are reported as adjusted p values (adj. p-value). Significance was defined as adj. p < 0.05 and |log2FC| ≥ 0.5.

To validate these findings, we performed RNA sequencing in cultured cell lines after knocking down KLF2 and CREB5. Consistent with *CellOracle* predictions, perturbation of the two hub TFs produced opposite fate biases: si*KLF2* versus siNC recapitulated an EndoMT/SMC-shift with release of endothelial homeostatic braking and suppression of adipogenic/lipid programs, whereas si*CREB5* versus siNC broadly repressed the endothelial activation-inflammation-adhesion axis and SMC/EndoMT effectors while upregulating metabolic modules supportive of adipogenesis (**Figure 7G-7H**). GSEA corroborated these directions, si*KLF2* enhanced the inflammation-adhesion-remodeling axis (complement, chemokines, collagen/ECM, blood microparticles) and suppressed translational/lipid-anabolic capacities (cytosolic ribosome, cytoplasmic translation, serine biosynthesis, long-chain fatty-acid binding), while siCREB5 showed negative enrichment of chemokine/cytokine signaling and leukocyte migration with positive enrichment of redox homeostasis, serine/one–carbon metabolism, iron-sulfur cluster assembly, and malonyl–CoA biosynthesis (**Supplementary Figure S6A-S6B**). Together, these results position KLF2 as a homeostatic brake whose loss drives EC fate toward SMC-like remodeling and, conversely, indicate that CREB5 knockdown inhibits the SMC branch while promoting adipogenic differentiation.

## Discussion

Sex differences in atherosclerosis have been recognized for decades, yet published studies have focused primarily on clinical investigations, and most basic research predominantly utilizing male animals ^10^. As the European Society of Cardiology (ESC) and the American Heart Association (AHA) advocate for reporting sex and gender in research ^82,83^, there is an urgent need to elucidate the mechanisms underlying sex-biased clinical manifestations of cardiovascular diseases.

EC monolayer dysfunction is a critical driver of atherosclerosis initiation and progression, a process known to be modulated by female sex hormones ^84,85^. While smooth muscle cells (SMCs) play a major role in plaque progression via phenotypic switching ^15,86^, the contribution of ECs to plaque cellular diversity remains understudied. Recent single-cell transcriptomic studies have provided an unprecedented view of the atherosclerotic plaque cell atlas ^87^. However, clinical samples derived from carotid endarterectomy (CEA) typically exclude the tunica media and adventitia, capturing only compositional snapshots rather than dynamic biological processes ^88^. Therefore, to capture the continuous dynamics of plaque initiation and progression, the use of animal models combined with genetic lineage tracing is indispensable.

In this study, we generated *Cdh5-CreER^T^*^2^/*Apoe^−/−^*/*Rosa26-mTmG* mice quantitatively trace endothelial dynamics in a hyperlipidemia model ^89^. Following tamoxifen-induced *Cre* activation, all ECs are labeled with GFP, which is subsequently inherited by all EC-derived cells as atherosclerosis progresses, allowing us to track EC cell fate and analyze the drivers of lineage commitment ^90^. Our analysis reveals that the contribution of EC transdifferentiation to atherosclerotic plaque formation may be substantially higher than previously estimated. A key finding of our study is the identification of a potential direct transdifferentiation route from ECs to adipocyte-like cells. Unlike the canonical Endothelial-to-Mesenchymal Transition (EndoMT) ^24^, our trajectory analysis indicates that this transition may occur without passing through a distinct mesenchymal intermediate. Our data provide direct evidence supporting the potential transdifferentiation of ECs into adipocyte-like cells and identify key transcription factors that may be involved in this process. These data provide direct transcriptomic evidence supporting EC plasticity toward an adipogenic fate and identify key transcription factors orchestrating this process. These findings challenge the conventional understanding of EC biology and propose a novel mechanism in the early pathogenesis of atherosclerosis. Future investigations comparing EC-derived adipocyte-like cells with SMC-derived adipocytes or foam cells will further clarify their distinct functional roles within the plaque.

We observed a significant sex preference in EC transdifferentiation. Female ECs exhibited a more dynamic phenotype, with a greater proportion transdifferentiating into SMC-like and fibroblast-like cells, a trajectory consistent with the clinical observation of thicker fibrous caps and more stable plaques in premenopausal women ^9^. In contrast, male ECs preferentially differentiated into macrophage-like cells, potentially contributing to plaque inflammation. By integrating snRNA-seq regulatory network analysis with bulk RNA-seq data, we demonstrated that estrogen modulates key nodes within the gene regulatory network (GRN), thereby influencing EC phenotypic plasticity. Furthermore, we identified and preliminarily validated key transcription factors (KLF2 and CREB5) that govern these distinct differentiation paths. Collectively, our findings highlight a sophisticated, sex-dependent regulatory program driving EC fate.

Lineage-tracing is a powerful tool to study the cell fate and commitment in a dynamic process. However, we acknowledge certain limitations inherent to the *Cdh5-CreER^T^*^2^ system. While this driver provides high-efficiency labeling, high *Cre* recombinase activity has been reported to induce cellular stress in some contexts ^91^. Although we observed a reduction in the total EC ratio in our reporter mice compared to non-Cre controls, this effect is likely systemic and does not invalidate the relative trajectory inference of the surviving, labeled GFP^+^ population. Our primary objective was to map the qualitative fate of ECs during disease progression, which remains robust despite these variations. Additionally, while we utilized the *Apoe*^−/−^ model, future studies in *Ldlr*^−/−^ mice or other atherosclerosis models would enhance translational relevance. Finally, incorporating spatial transcriptomics in future work will help to spatially resolve these sex-specific mechanisms within the complex plaque architecture.

In summary, this study delineates the distinct cell fates and trajectories of ECs during atherosclerosis progression in males and females. We identify key regulators driving these processes and propose a direct transdifferentiation route from ECs to adipocyte-like cells. These insights reshape our understanding of vascular plasticity and highlight the importance of sex-specific therapeutic strategies in atherosclerosis.

## Acknowledgments

This work was supported by grants from the National Natural Science Foundation of China (82270908, 82470464, 82570558) and the Shanghai Municipal Commission of Science and Technology (24ZR1445100).

## Conflict of interest

The authors declare no conflict of interest.

## Author Contributions

Q.R.L. and Y.Y.L. designed the study; C.L. performed all bioinformatics analysis; Y.Y.L. and S.M.W. performed animal experiments, cell biology, and immunostaining validation; C.L. drafted the manuscript; Q.R.L. and Y.Y.L. revised the manuscript. Q.R.L. and Y.Y.L. supervised the study.

## Data Availability Statement

The raw and processed data from the snRNA-seq and bulk-seq analyses are available from the corresponding author upon reasonable request.

## Supplementary Figures Legends

**Supplementary Figure S1.** Marker-based annotation of endothelial subclusters. **A.** Heatmap of representative marker genes across the nine endothelial subclusters, with side bars indicating EC type (arterial, capillary, lymphatic) and EC subtype. Expression is z–score normalized per gene. **B.** Augur perturbation analysis indicates differential perturbation in different EC subtypes using the area under the classifier curve (AUC); the dashed line equals 0.5. **C-K.** For each EC subtype (Art_EC-*Sox17*^+^, Lym_EC, Art_EC–*Mki67*^+^, Art_EC–*Myh6*^+^, Art_EC-*Fmo2*^+^, Art_EC-*Col8a1*^+^, Art_EC-*Ly6a*^+^, , Cap_EC, Art_EC-*Myh11*^+^): left, line plots showing average expression profiles of cluster markers across all EC subclusters; middle, violin plots of canonical markers within the indicated subtype; right, bar charts summarizing mean expression of additional markers.

**Supplementary Figure S2.** Sex-stratified differential expression and pathway enrichment across endothelial subtypes. **A-D.** Left: Volcano plots of DEGs (EFe vs EM) for Art_EC-*Myh11^+^* (A), Art_EC-*Ly6a^+^* (B), Cap_EC (C), and Lym_EC (D). Node size reflects STRING PPI degree (more connections = larger nodes); edges depict known interactions. Significantly up- and downregulated genes are colored as indicated. Middle: Semantic similarity maps summarizing GSEA results for each subtype. Circles represent GO terms positioned by semantic relatedness to reduce redundancy; circle size corresponds to term size, and color indicates enrichment direction. Right: Bar charts of selected GSEA terms for each subtype, reported as absolute normalized enrichment scores (NES) with direction (up/down) indicated by color. For volcano plot, differential expressions were computed using Wilcoxon rank-sum test. P values were adjusted for multiple testing using the Benjamini–Hochberg procedure to control the false discovery rate (FDR) and are reported as adjusted p values (adj. p). Significance was defined as adj. p < 0.05 and |log2FC| ≥ 0.5. For semantic similarity dot plot, semantic similarity-based clustering of GO terms from GSEA results. Each bubble represents the representative term of a GO cluster; positions reflect pairwise semantic similarity embedded to 2D using MDS (Semantic space X/Y). Colors encode log10(p-value) taken directly from the GSEA results, and the nominal p-value is permutation-based: for each gene set, ES is recomputed under permutations to form a null distribution, and the p-value is the proportion of null ES scores in the same direction with magnitude ≥ the observed ES (minimum ≈ 1/(number of permutations+1)).

**Supplementary Figure S3.** Sex-stratified differential expression and pathway enrichment across endothelial subtypes. **A-E.** Left: Volcano plots of DEGs (EFe vs EM) for Art_EC-*Col8a1^+^* (A), Art_EC-*Fmo2^+^* (B), Art_EC-*Sox17^+^* (C), Art_EC-*Mki67^+^* (D), and Art_EC-*Myh6^+^* (E). Node size reflects STRING PPI degree (more connections = larger nodes); edges depict known interactions. Significantly up- and down-regulated genes are colored as indicated. Middle: Semantic similarity maps summarizing GSEA results for each subtype. Circles represent GO terms positioned by semantic relatedness to reduce redundancy; circle size corresponds to term size, and color indicates enrichment direction. Right: Bar charts of selected GSEA terms for each subtype, reported as absolute normalized enrichment scores (NES) with direction (up/down) indicated by color. For volcano plot, differential expressions were computed using Wilcoxon rank-sum test. P values were adjusted for multiple testing using the Benjamini–Hochberg procedure to control the false discovery rate (FDR) and are reported as adjusted p values (adj. p). Significance was defined as adj. p < 0.05 and |log2FC| ≥ 0.5. For semantic similarity dot plot, semantic similarity-based clustering of GO terms from GSEA results. Each bubble represents the representative term of a GO cluster; positions reflect pairwise semantic similarity embedded to 2D using MDS (Semantic space X/Y). Colors encode log10(p-value) taken directly from the GSEA results, and the nominal p-value is permutation-based: for each gene set, ES is recomputed under permutations to form a null distribution, and the p-value is the proportion of null ES scores in the same direction with magnitude ≥ the observed ES (minimum ≈ 1/(number of permutations+1)).

**Supplementary Figure S4.** GeneTrajectory visualization and marker expression across EC-derived lineages. **A.** Dot plot of myofibroblast/SMC-skewed EndoMT markers across EC, Adipo-like, and SMC-like groups. Dot size indicates the fraction of expressing cells; color denotes mean expression. **B.** Dot plot of extracellular matrix remodeling and adhesion markers across the three groups, encoded as in panel C. **C.** Dot plot of adipocyte markers across EC, Adipo-like, and SMC-like groups, encoded as in panel C.

**Supplementary Figure S5.** DREMI/MI analysis of TF–gene associations. **A-I.** For each TF-target pair, four panels are shown: scatter plot of input expression values (left), kNN density estimate, joint probability heatmap with mutual information (MI), and conditional probability heatmap with DREMI score. Top rows display associations between Esrrg and selected adipocyte/brown-fat genes (*Ucp1*, *Ppargc1a*, *Ppargc1b*, *Cidea*, *Nrg4*, *Plin5*). Bottom rows display associations between Creb5 and ECM/lineage-related genes (*Col1a2*, *Fn1*, *Gata6*). Color scales indicate probability density.

**Supplementary Figure S6.** GSEA of bulk RNA-seq following TF knockdown. **A.** Representative enrichment plots for siKLF2 versus siNC. Each panel shows the running enrichment score, ranked gene list, and NES/P/FDR for selected GO terms. **B.** Representative enrichment plots for siCREB5 versus siNC. Each panel shows the running enrichment score, ranked gene list, and NES/P/FDR for selected GO terms.

## Notes

### Competing Interest Statement

The authors have declared no competing interest.

